# Staying on track: collagen fibers orientation strongly affects thrombi formation at high shear

**DOI:** 10.1101/2024.12.06.627297

**Authors:** E.A. Melnikova, P.H. Mangin, M.A. Panteleev, D.Y. Nechipurenko

## Abstract

**Objective:** Subendothelial matrix exhibits a distinctive organization, with collagen fibers beneath the endothelium oriented primarily parallel to the flow and comprising mainly types III and I collagen in arteries and arterioles. However, the significance of such organization in initiating rapid thrombus formation remains unclear.

**Approach and Results:** To investigate the role of collagen fibers orientation in triggering thrombus formation we utilized the *in vitro* microfluidic model of thrombosis. Human whole blood was perfused through a system with collagen fibers oriented either parallel or perpendicular to the flow, and the primary stages of platelet adhesion and thrombi growth were analyzed using a high-speed fluorescence microscopy.

At the shear rates of 200 and 1000 s^-1^ no significant difference in thrombogenicity was observed between the collagen fibers orientations. However, at a high shear rate of 2000 s^-1^, thrombi on the parallel fibers were higher and covered a much larger area than those on the perpendicular ones. High-speed microscopy revealed that platelets were able to achieve stable adhesion only after translocating for over several seconds under all studied conditions. Analysis of single platelets dynamics revealed that platelets interacted longer and translocated farther during their interaction with parallel fibers at high shear, suggesting that flow-aligned fibers facilitate stable adhesion by enabling platelets to translocate along them and stay in contact with collagen. Importantly, only a small fraction of collagen fibers belonging to type III collagen admixture - a common component of a standard type I collagen preparation - accumulated plasma vWF and supported platelet adhesion at high shear rate.

**Conclusion:** Type III, but not type I collagen fibers oriented parallel to the flow facilitate stable platelet adhesion at high shear rates. Thus, axial orientation of subendothelial collagen fibers observed *in vivo* may have physiological relevance for triggering rapid thrombus formation at high shear rates.

**Graphical abstract:** 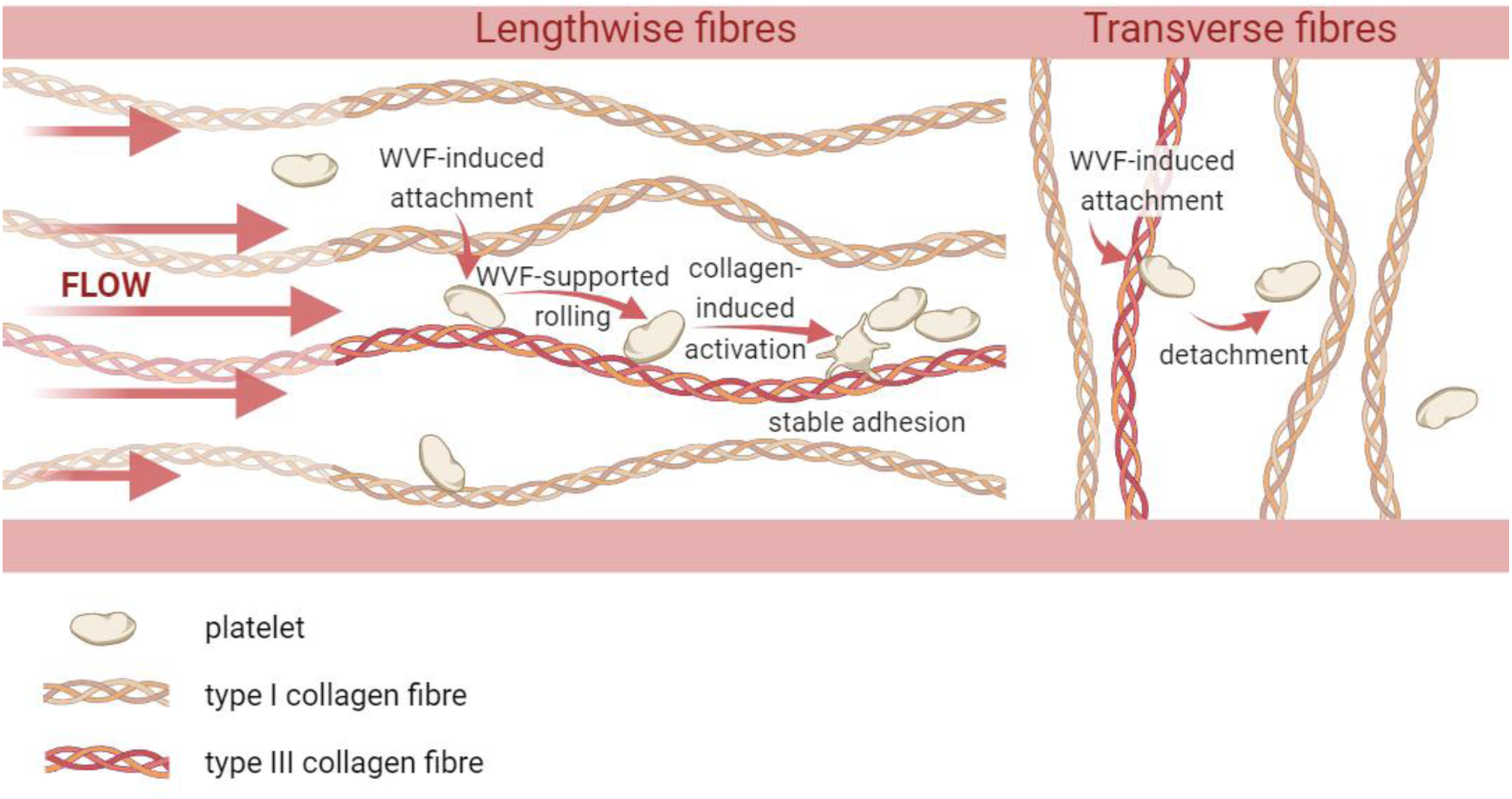

## Introduction

Understanding the molecular mechanisms regulating hemostasis in health and disease remains an acute fundamental task of physiology. Despite many decades of research, complications caused by impaired hemostatic response - bleeding and thrombosis - remain the leading causes of death and disability worldwide^1^. Specific interactions of the molecular and cellular blood components with subendothelial matrix proteins upon vessel wall injury play a central role in triggering primary hemostasis^2,3^. The biomechanical stage of primary hemostasis, namely the interaction between platelets and the vessel wall components, is currently considered as a key event that initiates platelet aggregate formation^4,5^. Analyzing how various components of the subendothelial matrix contribute to this process is significantly complicated by the vascular wall heterogeneity in terms of molecular composition and fibrillar components orientation^6^.

It is now known that vessel walls, especially arteries, contain a significant portion of fibrillar collagen which help to maintain their integrity and flexibility^7,8,9^. Moreover, collagen fibers in the vessel walls have preferential orientations: there is a thin layer of fibers aligned with the blood flow just below the endothelium; however, deeper into the arterial media, the orientation changes by 90° and the next thicker layer has fibers oriented perpendicular to the blood flow^10,11^. It has been suggested that such spatial organization might be important for countering both axial and circumferential mechanical stress^10^, however, the role of preferential orientations of collagen fibers in triggering primary hemostatic response upon superficial vessel wall injury remains unclear.

Here we take advantage of the microfluidics to analyze thrombus formation as a function of both collagen fibers orientation and the shear rate. Our results demonstrate that at a high shear rate fibers that are aligned with the flow are significantly more potent in triggering thrombus formation compared to the transverse fibers. We show that this effect is a consequence of primary platelet adhesion, which is much more efficient in case of lengthwise fibers due to primary platelet rolling. Our results suggest that axial orientation of collagen fibers in the subendothelium might be relevant for triggering rapid hemostatic response at high shear conditions and give insight into the optimal design of the microfluidic experiments that mimic hemostatic response upon vessel wall injury.

## Materials and methods

### Flow chambers preparation

The PDMS (SYLGARD 184 Silicone Elastomer) base was mixed with a curing agent at a 10:1 ratio and then cast onto a silicon master created using a standard photolithography. To define the dimensions of the PDMS structure, solid PDMS frames were used as borders to create a 4-5 mm high layer of PDMS. The plate was left for 5-10 minutes to allow air bubbles to escape and then incubated for 15 minutes at 150 ℃.The silicon master was designed to produce three rectangular channels, each measuring 1 cm in length, 1 mm in width, and 50 μm in height (see Figure 1). Following the incubation, the PDMS section containing the channels was carefully cut out from the borders using a scalpel. To facilitate the connection of tubing, holes were made at the ends of the channels using a 1 mm biopsy punch.

**Figure 1.**
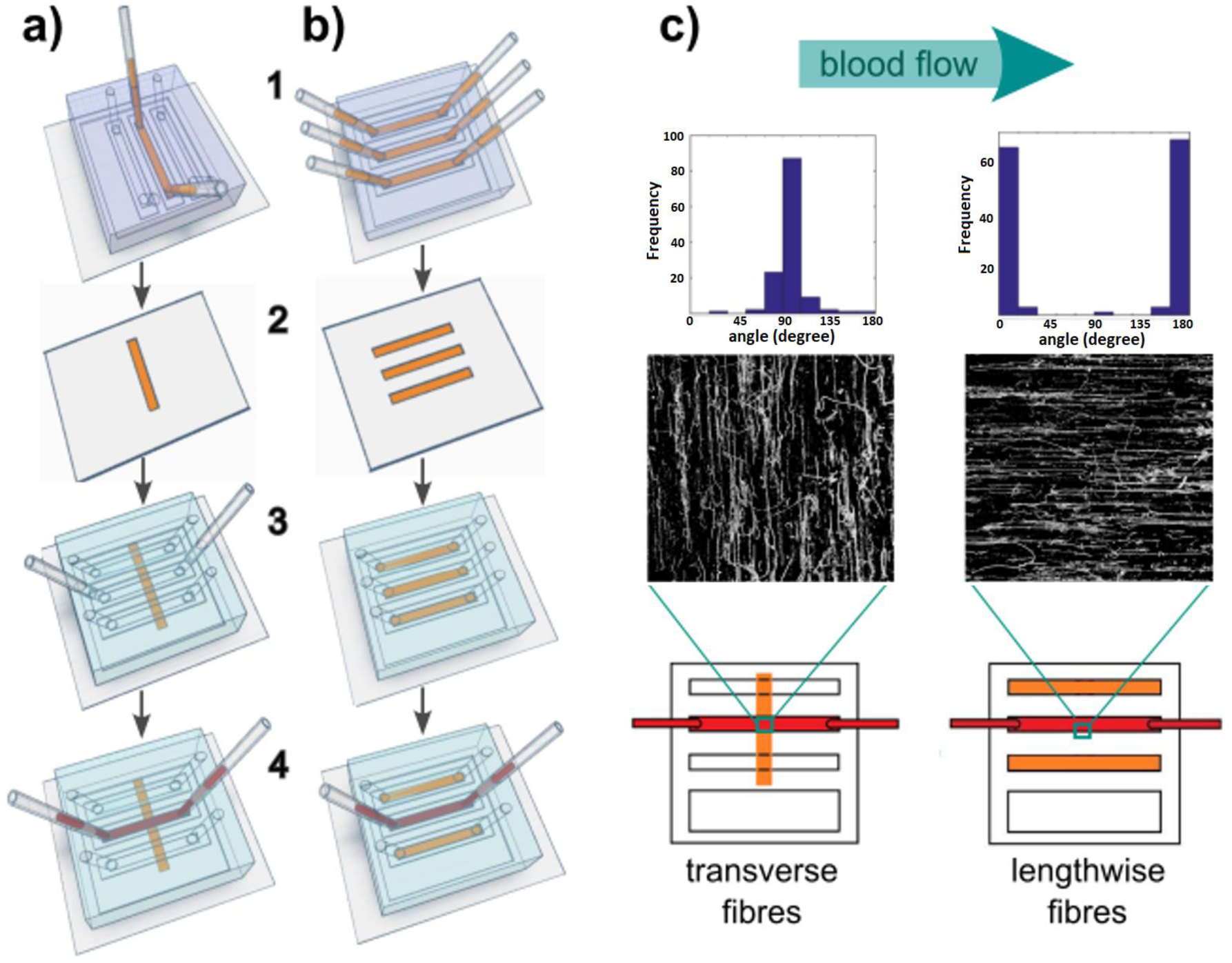
Schematics of the oriented collagen immobilization and blood perfusion. **a)**Transverse fibers preparation. 1. One channel of a PDMS chamber was incubated with a collagen solution (shown in orange) to apply the fibers to the glass coverslip. 2. The coverslip was dried on air. 3. The second chamber was mounted on the glass slide with immobilized collagen with channels oriented perpendicular to the collagen strip. 4. Blood was then perfused through one or several channels (shown in red). **b)** Lengthwise fibers preparation. 1. Three channels of a PDMS chamber were incubated with a collagen solution. 2. The coverslip was dried on air. 3. The second chamber was mounted on the glass slide with immobilized collagen, its channels were located at the same places as the channels of the first chamber used for collagen application. 4. Blood was then perfused through one or several channels. **c)** Collagen fibers, 40x magnification. Images were taken using bright-field microscopy, inverted and binarized with ImageJ and then analyzed using Curve Align software. The upper rows present orientation angle histograms along with corresponding binarized images of collagen fibers below; the lower row schematically depicts the orientation of the collagen strip (shown in orange) relative to the experimental chamber for obtaining transverse (left) and lengthwise (right) fibers (blood is shown in red).

### Blood collection

The study was conducted in accordance with the principles of the Declaration of Helsinki and under a protocol approved by the FRCC CHOI Ethical Committee.

Blood samples were obtained from healthy volunteers who confirmed that they had not taken any medication that could influence platelet function for at least two weeks prior to the phlebotomy. The first 2 ml of blood was discarded to ensure that the collected samples were free from any potential activators. Blood samples were collected using hirudin-coated tubes (S-Monovette® Hirudin, Sarstedt). Throughout the collection procedure, samples were gently mixed during blood withdrawal to ensure uniform hirudin dilution. Subsequently, the samples were stored at room temperature.

### Collagen immobilization

Glass coverslips were treated with low temperature air plasma and immediately covered with a PDMS flow chamber. Type I collagen solution from equine tendons (Chrono-Log Chrono-Par) was diluted in 20 mM acetic acid (1:4) before the application to get 0.2 mg/ml collagen solution. Collagen solution was injected into the channel(s) by placing a 20-200 μl pipette tip filled with the solution into the inlet hole 1 mm deep and gently pushing the plunger maintaining the same pressure for approximately 10 seconds until the channel and the outlet hole are filled as shown on Figure 1 (a, b). The chambers were incubated for 1 hour at room temperature in a humid environment. As we aimed to compare different collagen fibers orientation, there were two main methods of preparing collagen-coated slides. In order to create transverse fibers one channel of a chamber was filled with collagen solution via pipette (Figure 1a), for lengthwise fibers - all three of them (Figure 1b). For a number of experiments non-oriented fibers were obtained by placing a 10 μl drop of the collagen solution on a plasma-treated glass slide. Following the incubation, collagen solution was smoothly withdrawn from the channel(s) through the outlet hole with a pipette (for approximately 3 seconds), then the PDMS chamber was gently detached from the glass slide and discarded. Resulting collagen strip was rinsed with 300 μl of MQ by gently placing drops of liquid above the collagen-coated area on an angled slide using a pipette and letting them slide along the strip. Glass slides were then dried on air at RT.

Then, a new PDMS chamber was placed on the glass slide with immobilized collagen. The orientation of the chamber was adjusted based on the desired fiber orientation relative to the blood flow (as shown in Figure 1), because fibers are oriented along the channel that is used during their infusion. To passivate the surfaces and prepare the setup for further experiments, the channels were manually filled with an albumin-containing Tyrode buffer. This buffer included 150 mM NaCl, 2.7 mM KCl, 1 mM MgCl2, 0.4 mM NaH2PO4, 20 mM HEPES, 5 mM glucose, and 0.5% bovine serum albumin, with the pH adjusted to 7.4. This step was performed using a 2 ml syringe connected to the outlet tube.

### Mircrofluidic experiments

#### Thrombi growth dynamics

To label the platelets, we used DiOC6 (1 mM in DMSO), which was diluted to a concentration of 0.2 mM in PBS. This diluted DiOC6 solution was then added to either 200 or 400 μl of whole blood, along with 0.5 or 1 μl of DiOC6, respectively, in a test tube. To ensure proper mixing, the samples were gently mixed 15 times. Subsequently, the test tubes were incubated for 3 minutes. All these manipulations were carried out at room temperature. Labelled blood was perfused through a PDMS channel at varying shear rates: 200, 1000, or 2000 s^-1^. This was accomplished using a NE-1000 series syringe pump (New Era Pump Systems, Inc.) connected to the outlet tube, while the inlet tube was submerged in a separate test tube filled with blood (as illustrated in Supplementary Figure S1). During blood perfusion, z-stack images were captured at 40x magnification every minute, with a 2 μm step, to measure the height of the thrombi formed. At the end of each experiment, a photograph of the channel was taken at 4x magnification to measure the surface coverage of the formed thrombi. Microscopy setup is shown on the Supplementary Figure S1.

#### Dynamics of single platelet adhesion

Platelets were labelled as described earlier. Subsequently, the blood was perfused through a PDMS channel at shear rates of 1000 and 2000 s^-1^ for a total duration of 5 minutes, with all experiments conducted at room temperature. During the first 2 minutes of perfusion, high-speed timelapses were recorded in epifluorescence at either 40x or 100x magnification for individual platelet tracking. After the 5-minute perfusion, epifluorescence images were taken at 4x magnification to measure the surface coverage of the thrombi formed and to count the number of thrombi.

#### Collagen immunostaining

After the flow chambers were prepared following the previously described method, they were incubated with a 0.5% BSA (bovine serum albumin) solution. Subsequently, 10 μl of primary polyclonal rabbit antibodies specific to collagen type III (GeneTex) were introduced into the chamber using a pipette. The antibodies were mixed with PBS at a 1:200 ratio before each experiment. The chamber was then incubated for 30 minutes with antibody-containing solution. Following the primary antibody incubation, 10 μl of secondary goat anti-rabbit AlexaFluor647 (Invitrogen) antibodies were added to the chamber. These secondary antibodies are specific to the primary rabbit antibodies and enable the visualization of collagen type III binding sites. After an additional 30-minute incubation, polymer tubes were connected to the chamber, inlet tube was submerged into buffer A or PBS, outlet was connected to a syringe and antibodies were washed out by manually pulling the syringe plunger. With the antibody labelling process completed, bright-field and epifluorescence images were taken at 100x magnification. These images were used to compare and analyze the distribution of collagen fibers and collagen type III binding sites within the flow chamber. For the protocol on simultaneous vWF and type III collagen imaging see Supplementary materials (Page 1).

#### Interplay between VWF-collagen interaction and primary platelet adhesion

To visualize vWF, primary rabbit anti-human vWf antibodies were mixed with PBS (1:1000 ratio) and incubated with whole blood for 30 minutes. Then, secondary AlexaFluor568 goat anti-rabbit antibodies (mixed with PBS at 1:1000 or 1:200 ratio) were added for another 30 minutes. Meanwhile, bright-field images of collagen fibers were taken in the chosen field. Then platelets were labelled with DiOC6 and the blood was perfused through the PDMS channel at 2000 s^-1^ for 5 minutes. For the first 2 minutes of perfusion 15 fps timelapses were captured at 100x magnification in DiOC6 channel for the individual platelet tracking. After 2 minutes of perfusion, bright-field and epifluorescence images in the secondary antibodies channel were taken at 100x for quantification of collagen fibers stained for vWf, comparing them to all visible fibrils and studying their correspondence to platelet adhesion sites.

CurveAlign^12^ and ImageJ (National Institute of Health, Bethesda, MD, USA) software have been used for image processing, details can be found in Supplementary Materials (page 2).

### Computational modeling of primary adhesion

Stochastic model of primary platelet adhesion consisted of three modules that were implemented in Matlab. The first module generated random collagen fibers that were oriented either parallel of perpendicular to the blood flow. The second module randomly distributed vWF sites over the generated fibers, while the third module simulated platelet interactions with these fibers in the presence of flow. For details on simulation procedure and model parameters, see Supplementary Text 2.

## Results

### Lengthwise collagen fibers are more thrombogenic than transverse at high shear

To investigate how type I collagen fibers orientation affects thrombus formation, we performed *in vitro* experiments using hirudinated human whole blood and the widely used preparation of native collagen fibers (Chrono-par from Chronolog). In this study we used the microfluidic approach for collagen immobilization as well as for modelling the hemostatic response in an injured vessel. Applying collagen solution with a microfluidic chamber resulted in the alignment of collagen fibers with the flow, creating a strip of oriented collagen fibers on the coverslip’s surface, which served as the activator for thrombi formation (Figure 1 a, b). We used two orientations of collagen fibers: transverse (fibers perpendicular to the blood flow) and lengthwise (those oriented parallel to the flow). Fiber orientation was analyzed with a bright-field microscopy and CurveAlign software (Fig. 1 c). It is important to note that for all experiments collagen fibers were immobilized with the same method. The difference in the collagen fibers orientation was achieved by placing a perfusion chamber either parallel or perpendicular to the collagen strip (except for the special case of non-oriented fibers). Such chamber replacement procedure implies drying of a collagen strip, which could potentially affect its thrombogenicity. To address this concern, we compared our basic protocol for the lengthwise fibers with another one, which did not include replacement of a chamber (in this protocol the same chamber was used for both collagen application and blood perfusion, while collagen was always in contact with some solution). The results obtained for these 2 protocols were similar, so we concluded that chamber replacement and collagen drying did not impact the analyzed parameters (Supplementary Figure S2).

The experiments conducted at various flow rates demonstrated that thrombi growth on oriented collagen fibrils is influenced by the shear rate (Figure 2 (a-c)). We analyzed two parameters describing thrombi growth: mean thrombi height (mean h) and surface coverage (S). Notably, the effect of increasing shear rate from 200 s^-1^ to 2000 s^-1^ differed for lengthwise and transverse fibers. Mean height slightly increased with a shear rate on the lengthwise fibers and diminished on the transverse fibrils (Figure 2 b). Surface coverage exhibited more complex dependence on the shear rate: for the lengthwise fibers it increased insignificantly from 200 to 1000 s^-1^ and slightly diminished with further increase in shear rate to 2000 s^-1^. In contrast, the dependence for transverse fibers was completely different: while there was no significant difference in the surface coverage between 200 and 1000 s^-1^, there was a drastic decrease for 2000 s^-1^, leading to a significant difference between orientations at high shear (Figure 2 c). Intriguingly, thrombi growth dynamics on the non-oriented collagen fibers was similar to that on the transverse ones (Suppl. Figure S3).

**Figure 2.**
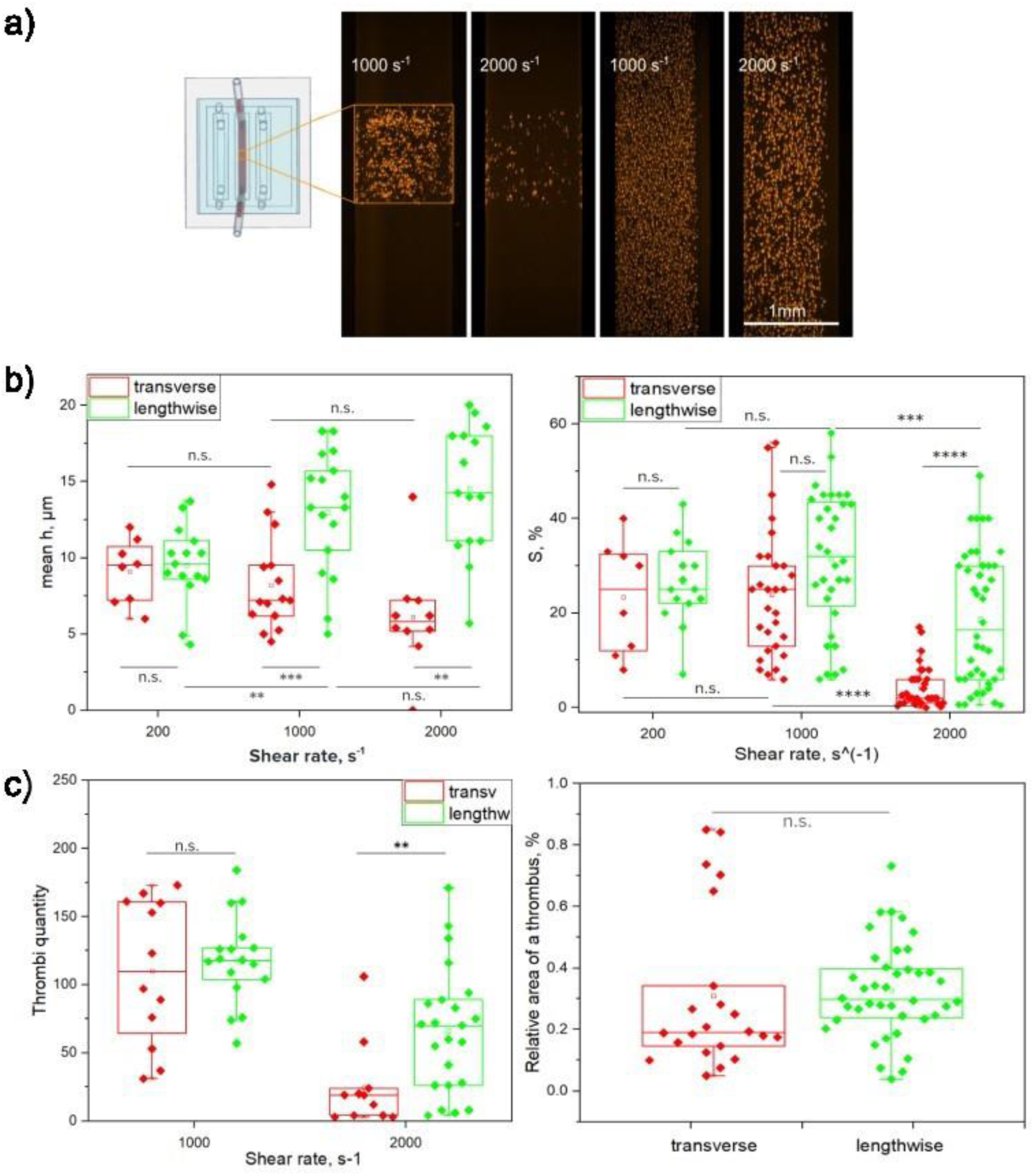
Effect of the shear rate and collagen fibrils orientation on thrombi parameters *in vitro*. **a)** Representative epifluorescent images of thrombi formed on collagen after 5 minutes of hirudinated whole blood perfusion. Schematic view of the PDMS chamber is given to indicate where the areas of observation were located. The images were taken right after 5 minutes of blood perfusion. Magnification - 4x; platelets were labelled with DiOC6. 1: Transverse fibers at 1000 s^-1^; 2: transverse fibers at 2000 s^-1^; 3: lengthwise fibers at 1000 s^-1^; 4: lengthwise fibers at 2000 s^-1^. **b)** (left) Mean thrombi height dependency on the surface shear rate. Red points correspond to the data acquired on the transverse collagen fibers; green - on the lengthwise. The boxes cover Q1-Q3 of the data, horizontal line shows the median. Data presented for N=15 experiments with 7 different donors. (right) Surface coverage (%) after 5 minutes of perfusion. N=35, 10 donors. Each data point corresponds to one run of the experiment (one channel). **c)** (left) Number of thrombi in the 330x330 μm^2^ area after 5 minutes of perfusion. (right) Relative area of a thrombus on the lengthwise and transverse fibers at 2000 s^-1^. The area was calculated as a total surface coverage in % divided by the number of thrombi. Each experiment consisted of the few runs (channels) and each run could have been carried out at a different shear rate or collagen fibers orientation using the blood of the same donor. Each point on a graph represents the data from one single run (i.e. single channel). Experiments carried out on the transverse fibers are marked in red color; those on the lengthwise fibers - in green. Middle line represents median value, upper and lower bounds contain data points in Q1-Q3 interval. Vertical axis corresponds to the value type, horizontal axis represents the wall shear rate. P-values of a Mann-Whitney test are shown for the corresponding data sets paired with a horizontal line: * p < 0.05; ** p < 0.01, *** p < 0.001; **** p < 0.0001.

To determine whether the observed effects were due to the difference in collagen stripe length (in the direction of the flow) for parallel and perpendicular orientations, we analyzed how local surface coverage varied along the channel. Оur results demonstrated that thrombi covered collagen uniformly (Supplementary Figure S4) on both orientations at 1000 s^-1^ and 2000 s^-1^, indicating that the observed phenomenon can not be attributed to the difference in the length of the activator-coated area.

As the surface coverage was significantly higher on lengthwise fibers compared to transverse fibers at 2000 s^-1^, we next focused on determining the mechanisms underlying the observed decrease in surface coverage. To get a deeper insight, we compared the quantity of thrombi formed on both orientations. This analysis revealed that at 1000 s^-1^, the number of thrombi formed on the transverse and lengthwise collagen fibers was similar. However, at 2000 s^-1^, the number of thrombi was significantly higher on the lengthwise fibrils (Figure 2c, left). Furthermore, the surface coverage to thrombi quantity ratio was similar for both orientations (p=0.1, Figure 2c, right), indicating that the difference in surface coverage at 2000 s^-1^ is primarily attributed to the difference in the number of thrombi and is likely associated with the initial stages of thrombi formation. To test this hypothesis, we aimed to analyze the primary stages of platelet adhesion to the oriented fibers.

### Platelets interact longer and translocate farther during their primary adhesion to the lengthwise fibers

To investigate the primary platelet adhesion at the high shear rate of 2000 s^-1^, we employed a high-speed fluorescent microscopy to track individual platelets and analyzed the critical parameters of their adhesion to the oriented collagen fibers. Analysis of the single platelet dynamics (Supplementary Video 1) revealed that many platelets attached to the surface, moved along it and eventually detached. These cells will be further referred to as “transient” platelets. However, a few cells were able to attach firmly - these will be further termed “stable” platelets. Importantly, the majority of platelets that managed to achieve stable adhesion, further initiated thrombus formation (Suppl. figure S5 (b)).

For hundreds of platelets, we measured the interaction time, defined as the period from the initial adhesion until the detachment in case of transient platelets and until the cessation of all movement in case of the stable platelets. We also determined their track lengths, representing the distance travelled by these platelets along the surface. Interestingly, evaluation of these parameters for stable platelets revealed that stable adhesion was typically preceded by rather lengthy translocation: both track lengths and interaction times of stable platelets were relatively high, spanning several microns and several seconds, respectively - significantly higher than those of transient platelets (Figure 3b). However, no significant difference was observed between collagen fibers orientations for both parameters, indicating that all stable platelets exhibited similar interaction times and track lengths regardless of collagen orientation at high shear. Expectedly, there were significantly more stable platelets on the lengthwise fibers than on the transverse ones (Suppl. Figure S5,a), consistent with the corresponding increase in thrombi quantity (Figure 2с). To investigate the origin of this difference, we next analyzed the dynamics of transient platelets. Both track lengths and interaction times for transient platelets were significantly shorter than those of stable ones (Figure 2b). This result suggests that to achieve stable adhesion and initiate thrombus formation, a platelet needs either to spend enough time travelling along the surface or travel a sufficiently long distance. Comparison of transient platelet track lengths and interaction times between orientations revealed that both parameters were significantly higher for lengthwise fibers compared to the transverse ones (Figure 3c). This indicates that at high shear rate, platelets adhering to lengthwise fibers generally translocate farther and interact longer with collagen than those rolling across the transverse fibers. As both the prolonged track length and translocation time appear to be prerequisites for stable adhesion (Figure 3b), these results could potentially explain the higher number of stably adherent platelets and, subsequently, more thrombi formed on the lengthwise fibers at 2000 s^-1^.

**Figure 3.**
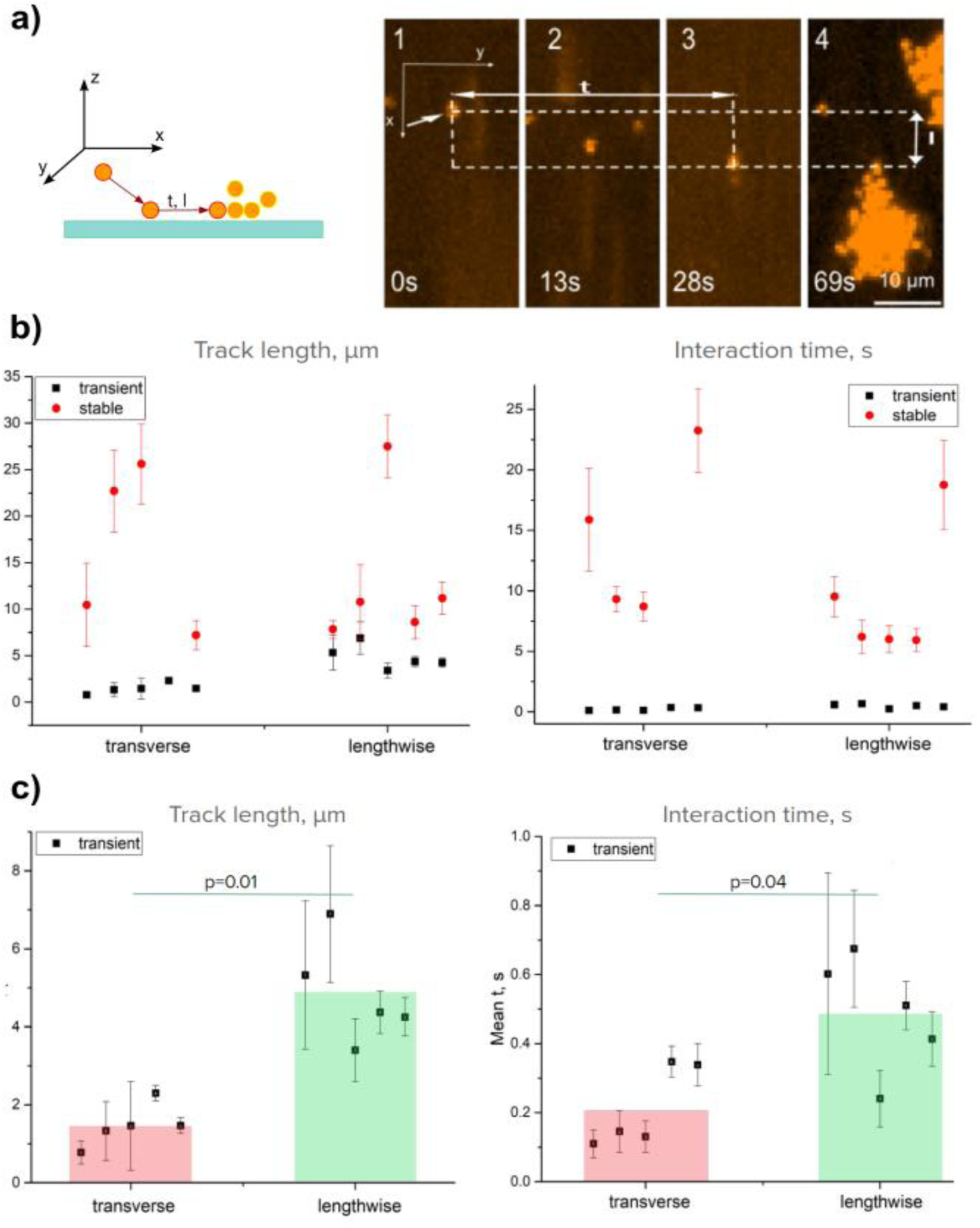
Analysis of single platelet attachment dynamics using a high-speed imaging. **a)** Schematics of a platelet attachment to the collagen-coated surface showing the critical studied parameters. Sequence of epifluorescence images from different time points of the experiments: 1) platelet attachment, 2) translocation, 3) stable adhesion, 4) thrombus formation. Track length is measured as a distance between platelet positions at 1) and 4), while interaction time - as an interval between 1) and 3).A platelet was considered stable if it ceased all movement and stayed in one place until the end of an experiment (and no less than 10 seconds). **b)** (left) Distance in micrometres (track length) travelled by transiently and stably adherent platelets on the transverse and lengthwise collagen fibrils at 2000 s^-1^. (right) Time in seconds (interaction time) that transiently and stably adherent platelets spent traversing along the collagen-coated surface at 2000 s^-1^. Each dot represents one experiment, each containing several runs. In each run as many platelets as possible were analyzed. Data points corresponding to stable platelets are red; to transient platelets - black. In some cases, small numbers of stably adherent platelets led to the increased variations. Experiments were conducted in pairs on both orientations, so each one on the transverse fibers has a pair on the lengthwise except for the forth one in which there were no stable platelets on the lengthwise fibers. **c)** Data on transient platelets from panel b. (left) Distance in micrometres (track length) travelled by transiently adherent platelets on transverse and lengthwise collagen fibrils at 2000 s^-1^. (right) Time transiently adherent platelets spent traversing along the collagen-coated surface at 2000 s^-1^.

To explain longer platelet interaction with lengthwise fibers at high shear, we proposed that the hydrodynamic drag force and fiber alignment with the flow enabled platelets to translocate along the fibers without losing contact with collagen. Hence, lengthwise collagen fibers might serve as rails for primary platelet translocation: through interaction with collagen-bound vWF molecules, platelets can roll for several micrometres (and several seconds) along the fiber, eventually becoming activated. To test this idea, we conducted experiments involving simultaneous localization of collagen-bound vWF and platelet adhesion.

### Platelets roll over the vWF-positive lengthwise fibers at high shear

The significance of the von Willebrand factor-GPIbα axis in platelet adhesion to collagen at high shear rates is well established. To find the mechanism underlying the differences in platelet interactions with transverse and lengthwise fibers, we turned to the analysis combining vWf staining and real time platelet adhesion dynamics. In these experiments, we simultaneously detected stained platelets and vWf multimers using a high magnification. Platelet free plasma or whole blood with primary vWf antibodies was perfused through the channels with transverse and lengthwise fibers, followed by the analysis of the staining using the secondary fluorescent antibodies. It provided us with the platelet adhesion dynamics while simultaneously mapping the distribution of vWF on the collagen fibers.

This analysis revealed that under the high shear rate conditions of 2000 s^-1^ platelets attach to vWf-positive fibers and rarely adhere where there is no vWF signal (Figure 4a), meaning that immunofluorescence staining strongly correlates with functional vWF molecules that support platelet adhesion and does not significantly interfere with it. On the lengthwise fibers platelet tracks could be rather long (Figure 4a(1-2)) and many tracks are overlapping with vWF-stained fibers for long distances (Figure 4b), implying that platelets are rolling along VWF-positive fibers. Platelet translocation along the vWF-positive lengthwise fibers could be seen on Suppl. video 2. It likely explains the increased interaction times and translocation lengths observed for platelets adhering to the lengthwise fibers (Figure 3d). Conversely, on transverse fibers, platelet tracks were mostly short (Figure 4a (3-4)), indicating brief attachment periods (Suppl. Video 3). Expectedly, there was also no long-distance overlapping of tracks with vWf signal on the transverse fibers (Figure 4b). Thus, collagen fibrils orientation significantly affects the primary adhesion process: platelets tend to translocate along lengthwise fibrils while showing brief attachment to the transverse fibers (Figure 4c, Suppl. Videos 2, 3). The platelets translocating along fibrils often pause without firm tethering, resuming movement after a brief period (Figure 4c, Suppl. Videos 2).

**Figure 4.**
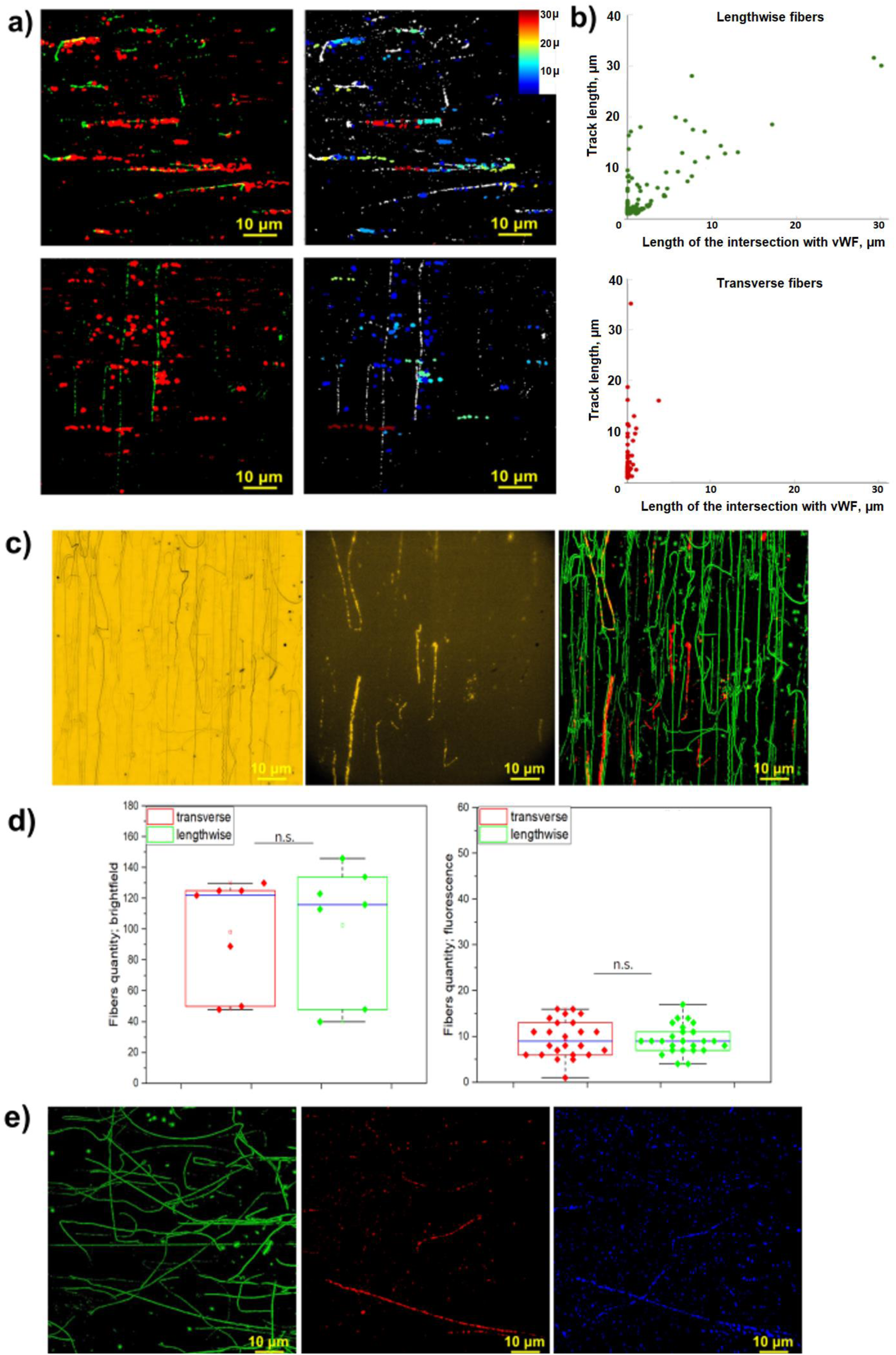
Analysis of the interplay between vWF-collagen interaction and primary platelet adhesion. **a)** Left column: representative images created by overlapping binarized epifluorescent images of the collagen-coated surface in the vWf antibody channel (green) and tracks of all platelets that interacted with the surface during 80 seconds of whole blood perfusion (red). Right column: the same images where collagen fibers are shown in white and platelet tracks in color according to the length of a track. The longer the track, the closer the color is to dark red. Color scheme is shown in the upper right corners of the images. **b)** Correlation between the total track length of the platelet and its intersection with VWF-positive part of the image for transverse (lower) and lengthwise (upper) fibers for the representative experiments (panel a). Each dot corresponds to a single platelet. **c)** Representative images of a collagen-coated surface in: bright-field (left), epifluorescence (von Willebrand factor antibody) (middle) channels and those binarized images overlapped (right; bright-field (all collagen fibrils) - green, epifluorescence (vWf) - red). Magnification: 40x. **d)** Number of fibers in 130x130 μm area in the bright-field (left) and epifluorescence (von Willebrand factor antibody) (right) channels for the transverse (red) and lengthwise (green) collagen orientation. **e)** Representative images of a collagen-coated surface in: (left) binarized bright-field, (middle) VWF antibody, (right) type III collagen antibody channels.

Interestingly, immunofluorescence analysis revealed that only a small portion of collagen fibers were stained with vWF antibodies. These vWF-positive fibers were generally thinner and barely (if at all) visible on the bright-field images (Figure 4c). Importantly, these findings suggest that only some of the immobilized collagen fibers had a high quantity of bound vWf molecules and, therefore, many GPIbα binding sites. Quantification of the images showed no significant difference in the intensity and overall quantity of stained fibers between collagen orientations (Fig.4 d). This implies that the differences in platelet adhesion between transverse and lengthwise collagen fibers cannot be explained by differences in vWf-collagen interaction. Consistent with our single platelet adhesion data, the number of platelets attempting to interact with the surface is similar for both collagen orientations (Suppl. Figure S6), supporting the idea that the total number of GPIbα binding sites should be comparable for both orientations.

We next proceeded to investigate the nature of these vWF-positive fibers in a “golden standard” of type I collagen preparations further. To test whether they belong to type III collagen admixture, we conducted immunofluorescent analysis, combining staining for adsorbed vWF and type III collagen antibodies (Figure 4a). To quantify the correlation between vWF and type III positive fibers, we developed an image analysis algorithm that utilized both bright-field and fluorescent images to integrate the signals along the visible fibers. The analysis revealed that approximately 80% of the fibers bearing a high fluorescent signal in either channel were simultaneously positive for both vWF and type III collagen. Moreover, platelet adhesion dynamics confirmed frequent platelet attachment to these fibers (Supplementary Figure S7). This strongly supports the conclusion that an admixture of type III collagen fibers is responsible for sustaining platelet adhesion at high shear rates.

### Stable adhesion requires prolonged interaction with collagen, leading to an increased thrombogenicity of the lengthwise fibers through the “rails” mechanism

To get a deeper insight into the mechanisms underlying the observed effects, we considered three hypotheses. The basic hypothesis suggests that in order to get activated and adhere stably, a platelet needs to sustain contact with collagen for a sufficient amount of time. In case of their interaction with the lengthwise fibers, platelets can translocate along vWF-rich collagen fibers during their primary adhesion and hence have more time for activation, resulting in a higher number of stably adhered platelets compared to the transverse fiber orientation. The second hypothesis suggests that a platelet needs to interact with a sufficient number of GPVI- activating collagen sites in order to achieve stable adhesion. In this scenario the lengthwise collagen fibers would be more thrombogenic due to the longer distances that platelets (on average) travel along such fibers during their primary adhesion thus interacting with a higher number of activating sites. Finally, according to the third hypothesis, in order to achieve stable adhesion, a platelet needs to encounter some “super-site” (e.g. a region with a high density of GPVI-activating sites on collagen). In this case, lengthwise fibers could be more thrombogenic due to seemingly higher probability for platelets to encounter such sites, because platelets translocate farther and hence “scan” more collagen surface during their primary interactions with such fibers.

Interestingly, experimental distributions of interaction times for transient platelets on both orientations (Figure 5 a) show a fast decline resulting in minimal numbers of platelets travelling more than 3-4 seconds. Importantly, the same distribution plotted for stable platelets peaked at 3-4 seconds (Figure 5 c), hence reinforcing the first hypothesis described above: as time-distributions plotted for two groups of platelets (transient and stable) have minimal overlap, the interaction time seems to be the main factor that allows platelet to become stable. Interestingly, the same analysis performed for platelet track lengths (Figure 5 b,d) showed a significant overlaps in track distributions for transient and stable platelets, meaning that it is unlikely that platelets need to travel far enough in order to become activated as suggested by the second hypothesis.

**Figure 5.**
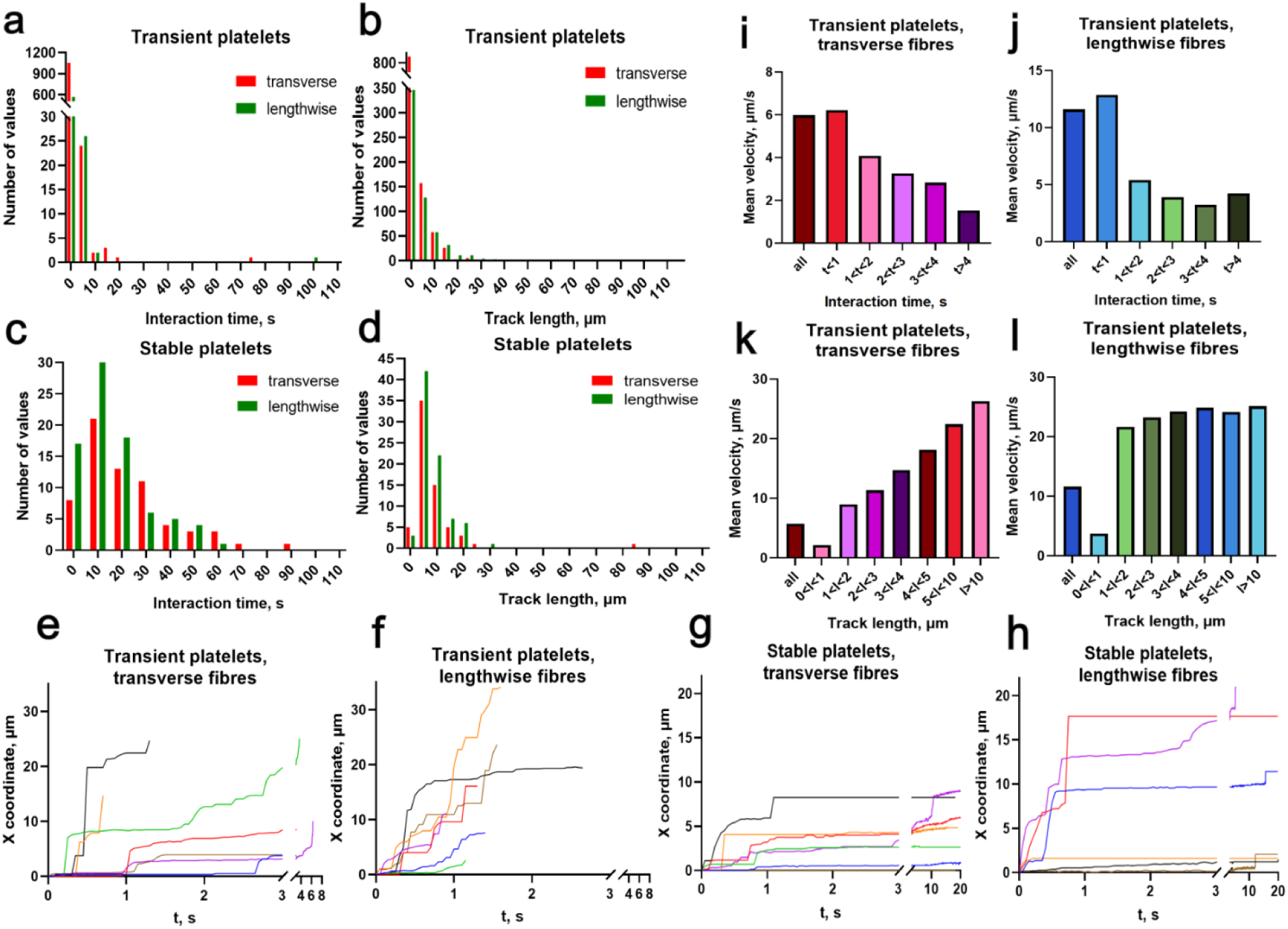
Analysis and computational modelling of individual platelet dynamics. a-d) Statistical distributions of individual platelet interaction times (a, c) and track lengths (b, d) for transient (a, b) and stable (c, d) platelets. Red bars correspond to the data acquired on the transverse fibers; green - on the lengthwise. N=10 experiments on each collagen orientation were analyzed. **e-h)** Individual platelet x coordinates (along the flow direction) from a representative experiment as the function of time. Each colored line corresponds to an individual cell. e, f: transient platelets, g, h: stable platelets interacting with: e, g: transverse fibers and f, h: lengthwise fibers. **i-l)** Velocity distributions for transient platelets as a function of total interaction time or total track length. The following binning intervals were chosen: 1 s for interaction times on transverse (i) and lengthwise (j) fibers and 1 μ for total track lengths on the transverse (k) and lengthwise (l) fibers.

Remarkably, analysis of the mean platelet velocities as a function of their interaction times performed for transient platelets showed a clear decrease in the average velocity with an increase of interaction time (Figure 5 i,j) for both collagen orientations, implying the time-dependent process of platelet activation after they start their interaction with collagen. Interestingly, analysis of the mean platelet velocities as a function of track length revealed a completely different picture (Figure 5k,l) with no decrease of the velocity with an increase of travelled distance. These data also reinforce the validity of the first hypothesis.

Detailed distance versus time plots for single transient platelets (Figure 5 e,f) revealed a complex nature of platelet movements: rather than moving with gradually decreasing continuous speed, they showed multiple accelerations and decelerations, leaped across some distances and slowed down on others. This stop-and-go character of platelet movement is characteristic for platelet-VWF interaction^13^. However, platelets rolling over the vWF-covered surface exhibit constant average velocity rather than clear decrease that we observed on collagen (Figure 5 e,f). The analysis of individual stable platelet tracks (Figure 5 g,h) confirmed higher platelet velocities in the initial stages of their translocation, while showing a significant decrease with time.

To get additional mechanistic insights, we turned to computational modelling of the primary platelet adhesion. We considered three types of models corresponding to three hypotheses described above. The physical core of all models was the same and considered stochastic interactions of platelets with vWF sites randomly distributed along collagen fibers (Figure 6a). Models were calibrated using the experimental distributions obtained for the transient platelets on the lengthwise fibers and then validated by inferring parameters of transient platelets on the transverse fibers and stable platelets on both fibers orientations and comparing it to the experimental data. The first model incorporated the general idea behind the basic hypothesis: in this model the probability of platelet detachment from vWF sites decreased with increase of total interaction time. In line with experimental data, some of the platelets slowed their movement with time (Figure 6b) and stayed on collagen for the whole simulation, so we considered them stable. This stochastic model adequately described experimental data for the transient platelets on the lengthwise fibers (Figure 6c,d) and predicted increased interaction times and track lengths for stable platelets on the lengthwise fibers (Figure 6e,f) (comparing to those of the transient platelets). The first model correctly predicted both the significant decrease of the number of stable platelets (Figure 6g) on the transverse fibers and the parameters of transient platelets on the transverse fibers compared to the lengthwise fibers, in line with the experimental data (Figure 6 c,d). The second model suggested that the probability of detachment from vWF sites decreased with increase of total number of vWF sites that the given platelet encountered during primary adhesion. This model quantitatively described experimental data for both transient and stable platelets on the lengthwise fibers, but failed to describe stable platelet adhesion to the transverse fibers. In case of the transverse fiber orientation, the total effective distance that platelets “scanned” throughout the fibers was too small compared to the lengthwise fibers, so no stable platelets were predicted by this model (Supplementary Figure S8a). The third model suggested that platelets can encounter some “super-sites” that lead to stable adhesion *in situ*. This model failed to describe experimental data on platelet adhesion even to the lengthwise fibers, as there was no difference between the parameters predicted for transient and stable platelets (Supplementary Figure S8 b).

**Figure 6.**
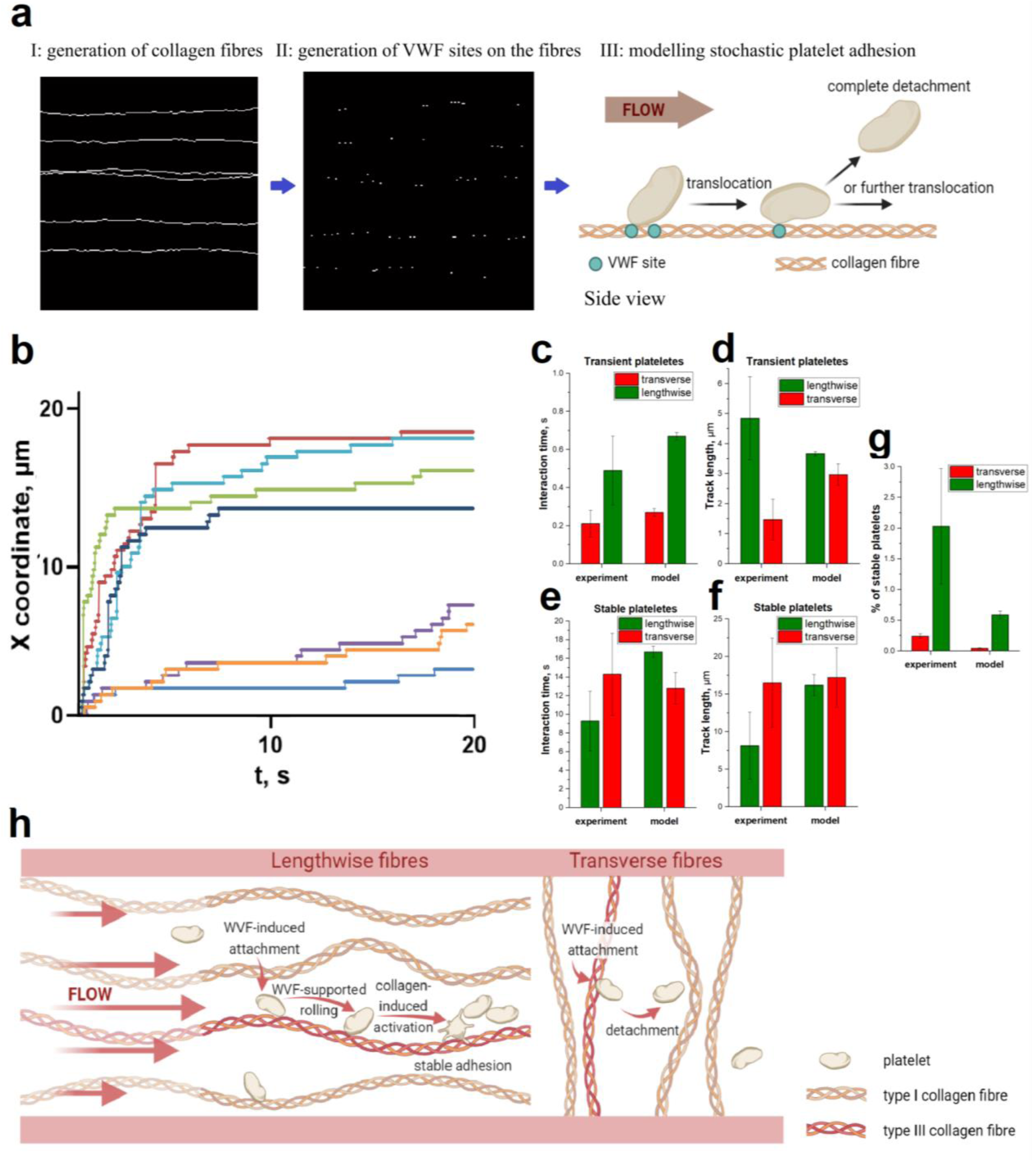
Computational modelling of primary platelet interaction with collagen. **a)** Schematics of the stochastic computational model of primary platelet adhesion to the oriented collagen fibers. Three basic stages of the modeling procedure are illustrated, for details see Supplementary Material. **b)** Individual platelet X coordinates (along the flow direction) as a function of time predicted by the model for the stable platelets in the lengthwise fibers. Each colored line corresponds to an individual platelet. **с-f)** Mean interaction time (left) and track length (right) of transient (c, d) and stable (e, f) platelets in the experiments (N=5 for each orientation) and the simulations (N=7 for each orientation) suggesting time-dependent decrease of platelet detachment from the vWF sites on collagen due to collagen-induced activation. Data for transverse fibers is shown in red; for lengthwise -in green. **g)** Mean stable platelets quantity as a percent of all attached cells for the experiments (N=5 for each orientation) and the simulations (N=7 for each orientation). Data for transverse fibers is shown in red; for lengthwise - in green.**h)** Schematics for the proposed mechanism of the increased thrombogenicity observed for the lengthwise collagen fibers at high shear. As platelets can roll over vWF-rich collagen fibers aligned with flow, they have more time for activation and hence have higher chances for stable adhesion, compared to the transverse fibers.

Taken together, the second and third models failed to describe the experimental data, reinforcing our conclusion that in order to adhere stably, a platelet needs to interact with collagen for a sufficient amount of time. Hence, the observed differences in thrombi formation on oriented fibers at high shear rate seem to arise from the combination of two main features: collagen-induced platelet activation takes time; platelets on the lengthwise fibers on average interact longer with collagen, as they can roll over the flow-aligned fibers during the primary vWF-dependent translocation (Figure 6h).

## Discussion

While many biochemical aspects of subedothelium matrix composition have been investigated in light of their contribution to the hemostatic response, several biomechanical features, including the role of internal mechanical stress and specific collagen fiber orientation in hemostasis, have not been thoroughly analyzed yet. In this work we used a microfluidic approach to investigate the role of native collagen fiber orientation in thrombus formation at various shear rates. Our results clearly indicated that the lengthwise collagen fibers are much more thrombogenic than the transverse fibers at high shear rates. While this finding might help to better understand the physiological relevance of axial collagen orientation right beneath the endothelial lining, it can be also relevant for designing more adequate microfluidic assays for both fundamental and clinical research.

Analysis of thrombus formation on native type I collagen fibers *in vitro* have been used for studying various aspects of the hemostatic response for several decades^14,5,15,16^. In these experiments, collagen can be immobilized on a surface in a variety of ways, leading to different results. Applying collagen solution in a drop leads to immobilization of non-oriented fibrils: while single fibers have some local orientation, there is no preferential orientation if many fibers are analyzed^17^ The use of the microfluidic flow chambers for collagen immobilization helps to acquire oriented fibrils on the coverslip surface as the fibers align with the direction of the flow during their infusion. The direction of these oriented fibers relative to the blood flow can be further controlled by the particular orientation of the channel used for further blood perfusion^18^. Interestingly, research teams around the world have obtained different results regarding the dependence of the collagen-induced thrombi formation on the shear rate. Some researchers have argued that thrombi size increases with shear rate, while some showed that there is a maximum at a certain value (as a rule, less than 1000 s^-1^) and smaller thrombi are formed with further increase of the wall shear rate (see Supplementary Table 1). Interestingly, analysis of the published protocols showed that in the first case mostly non-oriented or the lengthwise collagen fibrils were used, while in the second case the fibrils were oriented perpendicular to the blood flow. However, no direct evaluation of the impact of fibers orientation on thrombus formation has been performed.

In order to better understand the influence of the shear rate on thrombi growth and analyze the tendencies from different published studies, we combined the surface coverage data from the literature and plotted it on a single graph (Supplementary Figure S9). This generalized plot indicates that in case of the transverse fibers, surface coverage shows a decrease when wall shear rate is elevated higher than 1000 s^-1^ in all of the analyzed studies. Most of the experiments have shown the peak value for surface coverage on the transverse fibers at this shear rate. In contrast, surface coverage on the non-oriented and lengthwise fibers generally seems to increase with wall shear rate. Such difference in the dependence of thrombus formation on the shear rate for various orientations of fibers results in a significant difference of fiber thrombogenecity at high shear rate, in line with our findings. Interestingly, data points on the non-oriented and lengthwise fibers at 1500 s^-1^ lie very close to each other despite being taken from different studies. Moreover, surface coverage obtained at 1000 s^-1^ by different teams is quite similar although different experimental setups and reagents were used in these studies. Hence, if one is aiming at the application of thrombus formation on collagen *in vitro* as a clinical assay, 1000 s^-1^ might be the optimal wall shear rate for a robust system due to high reproducibility of the results. However, recent findings indicate that the rheology conditions at the early stages of hemostasis in case of a penetrating injury are characterized by the very high shear rates^19^ far exceeding the “standard” 1000 s^-1^. While the collagen fiber orientation within the injury site *in vivo* has not been investigated to date, our results suggest that in order to trigger rapid thrombus formation in case of the high shear rates, these fibers should be aligned with the flow. We propose that in order to accurately reproduce biomechanical aspects of primary hemostasis in response to penetrating injury, one should take into account both high shear rate and flow-aligned orientation of collagen fibers in the corresponding *in vitro* flow assays.

In line with the published data (Supplementary Figure S9), there was no significant difference between thrombogenicity of the lengthwise and transverse fibers at 1000 s^-1^ in our experiments. However, we observed a significant decrease in the thrombogenecity of the transverse fibers at 2000 s^-1^ – when primary adhesion of platelets to collagen relies on GPIbα – vWF interaction. In line with the recently published findings^20^, we directly observed that platelets exhibited high mobility before reaching stable adhesion on collagen.

Analysis of single platelet dynamics at the early stages of adhesion at high shear rates demonstrated that 1) the vast majority of interacting platelets experience only brief adhesion events and do not reach stable adhesion (transient platelets); 2) all platelets that achieved stable adhesion interacted with the surface for several seconds and translocated for several micrometers in case of both lengthwise and transverse fibers; 3) transient platelets on average interacted longer and translocated farther during their interaction with the lengthwise fibers. Further analysis revealed that it is platelet rolling over the vWF-covered single fibers that 1) explains generally longer interaction times of platelets with the lengthwise fibers; 2) precedes stable adhesion for the lengthwise fibers and 3) leads to the higher number of stably adhering platelets on the lengthwise fibers and hence higher thrombogenicity of these fibers at high shear.

Both thorough analysis of the experimental data and results of our simulations offer a simple explanation of the observed phenomenon: as platelet needs some time for activation and experiences vWF-dependent rolling before stable adhesion, the lengthwise fibers serve as rails that facilitate stable adhesion: platelets can roll along the fiber in the direction of the flow and not loose contact with collagen, thus having enough time for activation, in contrast to the transverse fibers. However, if this explanation works for the higher shear rates, then why there was no clear difference in thrombogeneicity of the lengthwise vs transverse fibers at 1000 s-1, wherein platelet adhesion also strongly depends on GPIbα – vWF interaction? We believe that the reason is the exponential decrease in the mean platelet “stop” time with an increase in shear rate reported for platelet rolling over the vWF surface^13,21^. According to the Bell law, there is an exponential increase in the molecular bond disassociation rate with the increase in the external drag force, which is proportional to the shear rate. Hence, the probability that some platelet will have a sufficient time for activation during it’s adhesion to the single vWF-covered transverse fiber before flow-induced detachment seems to be high enough at 1000 s^-1^, but too low for 2000 s^-1^. At 2000 s^-1^ the vast majority of platelets that attach to a single transverse fiber eventually detach before activation and completely loose contact with collagen, in contrast to the situation on the lengthwise fibers – wherein platelets can roll along the fiber.

Imunofluerescent analysis of vWF distribution over the collagen fibers revealed that only ≈10% of the visible fibers were stained with anti-vWF antibodies and supported both platelet adhesion and rolling at high shear. Moreover, these vWF-positive fibers were generally thinner and often barely visible in the bright-field. We next tested if these fibers belonged to type III collagen, as native collagens from tendons (Chrono-Par, Horm etc.) may contain admixtures of type III collagen. Simultaneous immunostaining for vWF and type III collagen confirmed this hypothesis. We also directly observed that type III collagen easily catches platelets at a high shear rate in our experiments (Supplementary video 4). These results are in line with the published data showing that type III collagen shows much higher affinity for vWF^22,23^, likely due to the completely conserved binding sequence, in contrast to the heterotrimeric nature of type I collagen. Importantly, our data clearly indicate that it is this relevant admixture of type III collagen that allows stable adhesion and thrombus formation at high shear rate conditions. Hence, our results imply that the pure type I collagen would be minimally thrombogenic at high shear due to minimal interaction of vWF with type I collagen fibers, challenging the general notion that type I collagen is a potent activator of thrombus formation at a wide range of the hemodynamic conditions^24^.

Taken together, our work shows that type III collagen fibers oriented parallel to the flow facilitate stable platelet adhesion at high shear rates because of the continuous interaction between this vWF-covered fibrils and platelets during their flow-directed translocation. Thus, axial orientation of subendothelial collagen fibers observed *in vivo* might be of a physiological relevance for triggering rapid hemostatic response at a high shear rate. Our findings also imply that both collagen immobilization procedure and type III collagen admixture in a “golden standard” preparation of type I collagen significantly influences the outcome of the microfluidic-based experiments and thus should be considered in their design and analysis.

## Author Contributions

E.A.M. developed the experimental model, carried out the experiments, analyzed the results and wrote the manuscript; P.H.M. analyzed the results and wrote the manuscript, M.A.P. designed the research, analyzed the results and wrote the manuscript, D.Y.N. outlined the scientific problem, designed the research, analyzed the results and wrote the manuscript.

## Acknowledgments

Authors thank Alexandr Ryabykh for the valuable discussions during the beginning of this work.

## Literature

1. Mensah, George A., Gregory A. Roth, and Valentin Fuster. "The global burden of cardiovascular diseases and risk factors: 2020 and beyond." Journal of the American College of Cardiology 74.20 (2019): 2529–2532.; Cardiovascular Diseases in Sub-Saharan Africa Compared to High-Income Countries: An Epidemiological Perspective, 2020

2. Ruggeri, Z. M., & Mendolicchio, G. L. (2007). Adhesion mechanisms in platelet function. Circulation research, 100(12), 1673–1685.

3. Brass, Lawrence F., and Scott L. Diamond. "Transport physics and biorheology in the setting of hemostasis and thrombosis." Journal of Thrombosis and Haemostasis 14.5 (2016): 906–917.; Flamm, Mathew H., and S. L. Diamond. "Multiscale systems biology and physics of thrombosis under flow." Annals of biomedical engineering 40 (2012): 2355–2364

4. Nieswandt, B., I. Pleines, and M. Bender. "Platelet adhesion and activation mechanisms in arterial thrombosis and ischaemic stroke." Journal of thrombosis and haemostasis 9 (2011): 92–104.;

5. Cosemans, Judith MEM, et al. "The effects of arterial flow on platelet activation, thrombus growth, and stabilization." Cardiovascular research 99.2 (2013): 342–352.)

6. van Geffen, Johanna P., et al. "High-throughput elucidation of thrombus formation reveals sources of platelet function variability." Haematologica 104.6 (2019): 1256.

7. Plenz GA, Deng MC, Robenek H, Völker W. Vascular collagens: spotlight on the role of type VIII collagen in atherogenesis. Atherosclerosis. 2003 Jan;166(1):1–11. doi: 10.1016/s0021-9150(01)00766-3. PMID: 12482545.

8. Barnes, M. J. (2018). Collagens of normal and diseased blood vessel wall. In Collagen (pp. 275-290). CRC Press.

9. Chow, M. J., Turcotte, R., Lin, C. P., & Zhang, Y. (2014). Arterial extracellular matrix: a mechanobiological study of the contributions and interactions of elastin and collagen. Biophysical journal, 106(12), 2684–2692.

10. Timmins, L. H., Wu, Q., Yeh, A. T., Moore Jr, J. E., & Greenwald, S. E. (2010). Structural inhomogeneity and fiber orientation in the inner arterial media. American Journal of Physiology-Heart and Circulatory Physiology, 298(5), H1537–H1545.

11. Schriefl, Andreas J., et al. "Determination of the layer-specific distributed collagen fiber orientations in human thoracic and abdominal aortas and common iliac arteries." Journal of the Royal Society Interface 9.71 (2012): 1275–1286.

12. Liu, Y., Keikhosravi, A., Mehta, G. S., Drifka, C. R., & Eliceiri, K. W. (2017). Methods for quantifying fibrillar collagen alignment. Fibrosis: methods and protocols, 429–451.

13. Coburn, L. A., et al. "GPIbα-vWF rolling under shear stress shows differences between type 2B and 2M von Willebrand disease." Biophysical journal 100.2 (2011): 304–312.

14. Neeves, K. B., Onasoga, A. A., Hansen, R. R., Lilly, J. J., Venckunaite, D., Sumner, M. B., … & Di Paola, J. A. (2013). Sources of variability in platelet accumulation on type 1 fibrillar collagen in microfluidic flow assays. PloS one, 8(1), e54680.

15. De Witt, S. M., Swieringa, F., Cavill, R., Lamers, M. M., Van Kruchten, R., Mastenbroek, T., … & Cosemans, J. M. (2014). Identification of platelet function defects by multi-parameter assessment of thrombus formation. Nature communications, 5(1), 4257.

16. Muthard, Ryan W., and Scott L. Diamond. "Side view thrombosis microfluidic device with controllable wall shear rate and transthrombus pressure gradient." Lab on a Chip 13.10 (2013): 1883–1891.

17. Hosokawa, Kazuya, et al. "A microchip flow-chamber system for quantitative assessment of the platelet thrombus formation process." Microvascular research 83.2 (2012): 154–161.

18. Neeves, K. B., et al. "Microfluidic focal thrombosis model for measuring murine platelet deposition and stability: PAR4 signaling enhances shear-resistance of platelet aggregates." Journal of Thrombosis and Haemostasis 6.12 (2008): 2193–220

19. Yakusheva, A. A., Butov, K. R., Bykov, G. A., Závodszky, G., Eckly, A., Ataullakhanov, F. I., … & Mangin, P. H. (2022). Traumatic vessel injuries initiating hemostasis generate high shear conditions. Blood advances, 6(16), 4834–4846.

20. Pugh, N., Maddox, B. D., Bihan, D., Taylor, K. A., Mahaut-Smith, M. P., & Farndale, R. W. (2017). Differential integrin activity mediated by platelet collagen receptor engagement under flow conditions. Thrombosis and haemostasis, 117(08), 1588–1600.

21. Kaneva, V. N., Dunster, J. L., Volpert, V., Ataullahanov, F., Panteleev, M. A., & Nechipurenko, D. Y. (2021). Modeling thrombus shell: linking adhesion receptor properties and macroscopic dynamics. Biophysical journal, 120(2), 334–351.

22. Brondijk, T. H. C., Bihan, D., Farndale, R. W., & Huizinga, E. G. (2012). Implications for collagen I chain registry from the structure of the collagen von Willebrand factor A3 domain complex. Proceedings of the National Academy of Sciences, 109(14), 5253–5258.

23. Moroi, M., and S. M. Jung. "A mechanism to safeguard platelet adhesion under high shear flow: von Willebrand factor–glycoprotein Ib and integrin α2β1–collagen interactions make complementary, collagen-type-specific contributions to adhesion." Journal of Thrombosis and Haemostasis 5.4 (2007): 797–803.

24. Savage, B., Almus-Jacobs, F., & Ruggeri, Z. M. (1998). Specific synergy of multiple substrate–receptor interactions in platelet thrombus formation under flow. Cell, 94(5), 657–666

